# Ptolemaea: consensus, comprehensive annotation of antiviral defence systems in bacterial genomes

**DOI:** 10.64898/2026.06.26.734901

**Authors:** Emmet B. T. Campbell, Timofey Skvortsov, Christopher J. Creevey

## Abstract

**Motivation:** Bacteria carry a large repertoire of antiviral defence systems, our knowledge of which is expanding rapidly. Several bioinformatics tools now exist to identify them. Though powerful, these tools can differ in the models they use and the nomenclature they return, thus a single tool could both miss an annotation and disagree with its peers.

**Results:** Here we describe Ptolemaea, a pipeline for harmonising phage-defence annotations across multiple tools by reconciling PADLOC, DefenseFinder, and a bidirectional BLAST. Over a common predicted set of proteins, Ptolemaea provides a consensus annotation list per genome. The pipeline is not intended to outperform or replace its component tools; its purpose is to maximise the number of defence systems recovered from a genome and to make disagreements between tools explicit and resolvable. We demonstrate the pipeline on 700 complete genomes spanning the ESKAPE pathogens and Escherichia coli, recovering 32,509 defence annotations, of which 50.6% were supported by more than one annotation source.

**Availability:** The Ptolemaea pipeline is freely available at https://github.com/ecampbell50/Ptolemaea.

**Supplementary information:** Genome accessions used in this analysis can be found in S1, and script for genome retrieval in S2. Collated consensus annotation counts can be found in S3, while all raw tool outputs and curated decisions for each species can be found in S4-10.

## Background

The rapidly expanding and diverse repertoire of mobile genetic element (MGE)-excluding defence systems in prokaryotes continues to be a fascinating and important area of research. Not only has the discovery of many of these systems led to the development of a number of biotechnological tools, but understanding their complexity is crucial for creating truly effective phage-therapies [1]. The rapid discovery of these systems is often attributed to the tendency of defence genes to cluster together in ‘defence islands’ [2], which can be mined for novel systems using a combination of guilt-by-association and homology-based approaches [3, 4, 5]. The detection of already known and characterised defence systems is thus central to the computational discovery of new ones.

Several tools now exist that allow the detection and characterisation of defence systems present in a genome of interest, notably PADLOC [4] and DefenseFinder [5], which use curated hidden Markov model (HMM) libraries together with gene presence/absence and synteny criteria to annotate complete systems. DefensePredictor, a recently published tool, takes an alternative route by using the protein embedding space of known genes to predict antiphage defences, based on a training set of both defence and non-defence genes [6].

For PADLOC and DefenseFinder, the annotation capability is limited by their underlying databases, which differ in content and are updated on different schedules. Also, an annotation assigned by one tool may be missed by another. Or where both tools provide an annotation for the same locus, either the nomenclature or biology of that annotation may differ. These issues are well illustrated by recent work in the *Bacillus cereus* group, where PADLOC and DefenseFinder were run side by side across 6,354 assemblies and showed both broad agreement and systematic differences in the systems they reported [7]. A consistent classification of defence systems is desired for any downstream analyses, to ensure reproducibility and to enable direct comparison between studies.

To meet this need we developed Ptolemaea, a pipeline that runs PADLOC, DefenseFinder and a homology-based BLAST search over a common, validated protein set, and merges their results into one consensus annotation per genome. Ptolemaea helps to maximise the number of defence systems recovered from a genome, and to make disagreements between tools explicit and resolvable. Below we describe the implementation and demonstrate the pipeline on a panel of 700 complete genomes spanning *Escherichia coli* and the ESKAPE pathogens (*Enterococcus faecium, Staphylococcus aureus, Klebsiella pneumoniae, Acinetobacter baumannii, Pseudomonas aeruginosa*, and *Enterobacter spp*.).

## Implementation

### Overview

Ptolemaea is a Singularity container command-line pipeline written in Bash and Python that takes one or more bacterial genome assemblies (nucleotide FASTA) as input and produces, for each genome, a reconciled table of predicted antiviral defence systems. It is structured as five sequential stages (Figure 1A):

**Figure 1.**
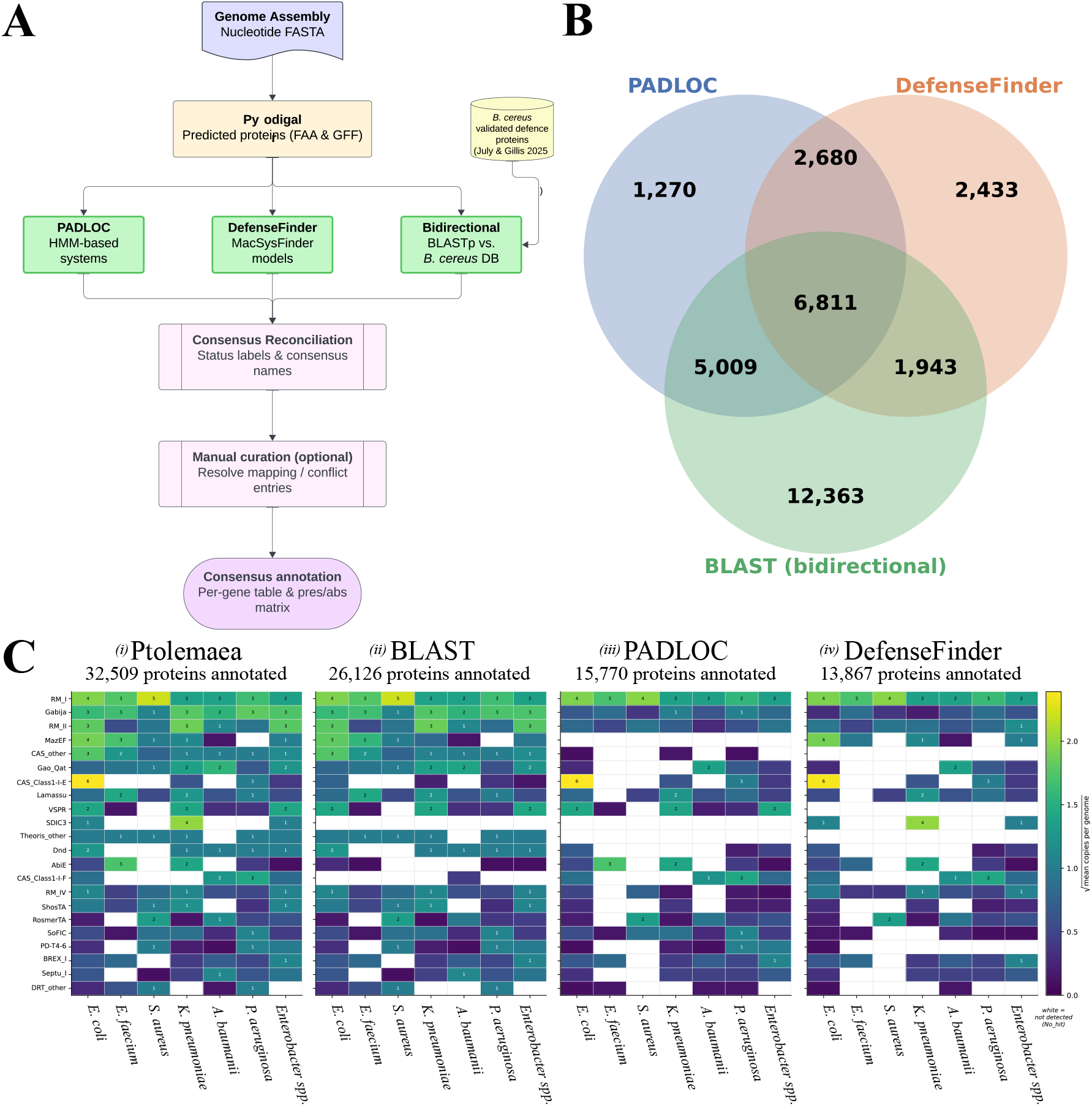
**A**: Ptolemaea pipeline overview. **B**: Venn diagram of each tool’s hits across the genome set. Overlaps show instances where tools provided an annotation for any given protein, regardless of what that annotation was. BLAST figures show only those that passed all homology filters. **C**: Defence-system landscape across the demonstration panel (700 genomes) and each annotation source, showing what each would return if used in isolation: *(i)*: the full Ptolemaea consensus, *(ii)*: BLASTp bidirectional, filter-passed hits only, *(iii)*: PADLOC hits only, and *(iv)*: DefenseFinder hits only. Per-genome copy number of the 22 most abundant consensus system subtypes (rows, ordered by overall abundance) across the seven species; colour is the square root of the mean copy number, and prominent cells (*≥*1 copy per genome) are annotated. A protein counts as missed when the tool’s raw output is No hit where another tool detected it; cells are drawn white where no protein of a subtype was detected in a species (mean = 0). Row label corresponds to the final consensus annotation, regardless of what each tool assigned.

1. **Gene prediction and protein extraction** Each input assembly is annotated with Pyrodigal [8] (v3.7.1) to produce predicted protein sequences (`.faa`) and the corresponding feature coordinates (`.gff`). Using a single, common protein set for all downstream tools ensures that every annotation source operates on identical locus tags, which is essential for meaningful comparison.
2. **PADLOC annotation**. PADLOC [4, 9] (v2.0.0; DB v2.0.0) is run on the Pyrodigal protein/GFF pair.
3. **DefenseFinder annotation**. DefenseFinder [5, 10] (DefenseFinder v2.0.1; MacSyFinder v2.0; CasFinder v3.1.0; hmmer v3.4), built on MacSyFinder v2 [11], is run on the same protein set.
4. **Bidirectional BLASTp against a curated defence-protein database**. BLASTp [12, 13] (v2.15.0) is performed in two directions against a database of validated defence-system proteins from the *B. cereus* group catalogue of July and Gillis [7]. In the forward search, predicted proteins from the query genome are used as queries against the *B. cereus* database. In the reverse search, the *B. cereus* proteins are used as queries against a temporary database built from the query genome’s predicted proteins. Both searches use a single-hit threshold (`--max target seqs 1`) and an E-value cut-off of 1 × 10^−5^. The two sets of results are subsequently evaluated together during consensus reconciliation to identify BLAST-only proteins with bidirectional support (see: Consensus logic: BLAST).
5. **Consensus reconciliation**. For each predicted protein that receives at least one annotation, the hits from the three sources are collated, a consensus annotation is assigned, and a status label records how the annotation was reached (see: Consensus logic). Proteins assigned a mapping or conflict status are flagged and retrievable using an auxiliary script so the user can fill in and re-submit to generate a fully resolved annotation.

For contemporaneous clarity, software versions have been provided, but these are subject to change as the underlying tools and databases evolve. Ptolemaea is designed to be flexible and modular, so that individual stages can be re-run with updated resources, with the potential to even add other tools.

### Consensus logic

#### Nomenclature harmonisation

A central challenge in combining PADLOC and DefenseFinder output is that the two tools use different naming conventions for the same defence systems. Ptolemaea resolves this via the master toolkey.csv, a look-up table derived from the unified nomenclature proposed by July and Gillis [7], which maps each tool’s system names to a shared consensus subtype name. For example, PADLOC’s ‘AbiD’ and DefenseFinder’s ‘Abi2’ are both mapped to the consensus name ‘Abi2DF’. Systems that are catalogued by PADLOC or DefenseFinder but were not validated in the *B. cereus* group reference set lack an entry in the master toolkey.csv and cannot be automatically assigned a consensus name; these require user input (see mapping below).

#### Status labels

Each predicted protein is assigned one of the following status labels, which are propagated to the output table and serve as an audit trail for every consensus decision:

##### agree

Both PADLOC and DefenseFinder detected the protein, and both map to the same consensus name in the master toolkey.csv. The consensus name is assigned automatically.

##### resolved

PADLOC and DefenseFinder detected the protein but mapped to different consensus names. The disagreement is settled by BLAST voting: each tool starts with one vote, and a forward or reverse BLAST hit whose annotation matches a tool’s consensus name awards that tool one additional vote (giving a maximum of three votes per tool). The tool with the majority vote is selected as the consensus annotation. These are assigned automatically.

##### single

Only one HMM-based tool detected the protein, and the system it detected has a consensus name in the master toolkey.csv. The single tool’s consensus name is assigned automatically and flagged as single-source support.

##### BLAST

Neither PADLOC nor DefenseFinder detected the protein, but BLASTp evidence from both search directions is present. A BLAST-only hit is accepted only when all four criteria are satisfied: both forward and reverse searches returned a hit; (ii) both hits agree on the system annotation; (iii) the ratio of query length to subject length from the forward hit satisfies 0.8 ≤ *Q/S* ≤ 1.25; and (iv) the alignment coverage from the forward hit, defined as 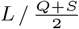, satisfies 0.8 ≤ cov ≤ 1.25, where *L* is alignment length, *Q* is query protein length and *S* is subject protein length. Criteria (iii) and (iv) guard against spurious hits between proteins of very different sizes or where only a short region aligns. Proteins that fail any of these criteria are discarded and do not appear in the output.

##### mapping

A protein was detected by one or both HMM-based tools, but the annotated system is absent from the master toolkey.csv. That is, the system was not validated in the *B. cereus* group reference catalogue and therefore has no agreed consensus name. No automatic assignment is made; these entries require the user to inspect the raw tool calls and supply a final annotation.

##### conflict

Both tools detected the protein, both have mappings in the master toolkey.csv, but BLAST voting resulted in a tie. The competing annotations are retained verbatim and presented to the user for adjudication.

Mapping and conflict entries are efficiently handled by running ‘extract unresolved patterns.py’ after the main pipeline. This script scans all per-genome consensus files, groups the flagged proteins by their unique combination of PADLOC, DefenseFinder and BLAST annotations, and writes a compact CSV template in which each unique pattern appears only once regardless of how many genomes or proteins share it. The user fills in the ‘TYPE’, ‘SUBTYPE’ and ‘OUTCOME’ columns for each pattern; those resolutions are then applied back to every affected protein in a single pass.

## Demonstration

### Genome dataset

To demonstrate the pipeline we assembled a panel of 700 publicly available complete genome assemblies spanning the ESKAPE pathogens together with *Escherichia coli*. Each species had 100 complete assemblies. Complete genome assemblies were retrieved from NCBI RefSeq (accessed: 05 JUN 2026) using the NCBI Datasets command-line tool ‘datasets v18.29.1’. All available complete assemblies were filtered using checkm-completeness ≥99% and checkm-contamination ≤2%, then ranked by release date (most recent first) and contig N50 (largest first); the top 100 assemblies per species were selected. An exception was made for *S. aureus*, for which no assemblies in RefSeq carried a completeness score ≥99%; a relaxed threshold of ≥98.85% was therefore applied, yielding a pool of 113 assemblies from which the top 100 were selected. Accession numbers for all assemblies are listed in Supplementary Table S1. Genome-retrieval script can be found in Supplementary S2. This panel was chosen to span clinically important taxa with well-studied but differing defence repertoires, not to test any specific hypothesis.

### Defence gene recovery

Ptolemaea recovered 32,509 defence-system annotations spanning 186 distinct named system types (Figure 1; Supplementary Table S3), where “named” excludes a catch-all ‘Other’ category and calls that remained unresolved after curation. The number of defence-system annotations per genome ranged from 17 to 103 (median 45) and varied markedly by species (Figure 1C). Recomputing the same landscape from the PADLOC or DefenseFinder calls in isolation recovered roughly half as many systems per genome (median 20 and 18 genes per genome, respectively, versus 45 for the full consensus) and left many system-type rows sparsely populated (Figure 1).

BLAST-only calls accounted for 12,363 hits, which were not returned by either PADLOC or DefenseFinder.

These represent homologs of the validated *B. cereus* defence proteins that passed the strict filters accounting for coverage and alignment.

Of the 32,509 consensus calls, 15.7% carried an agree status, 4.5% were resolved, 22.6% were single-source, and 38.0% were BLAST-only hits. The remaining 19.1% were flagged for manual curation (mapping, 18.3%; conflict, 0.8%) and resolved using the unresolved-pattern workflow. Just over half of all consensus calls (50.6%) were supported by two or more independent sources.

Crucially, no single source was sufficient on its own. Taking the union of all three sources recovered 32,509 annotations, roughly twice the number recovered by either HMM-based tool alone (PADLOC, 15,770; DefenseFinder, 13,867). Of the BLAST evidence, 13,763 annotations (52.7%) were corroborated by at least one HMM-based tool.

## Significance

We created a pipeline, Ptolemaea, that maximises defence system annotations in bacterial genomes. PADLOC or DefenseFinder may return proteins the other tool has missed, and may disagree or differ in the annotations they provide. Ptolemaea executes both tools, and automates and/or streamlines the process of outputting a consensus name between the two tools through use of an additional bidirectional BLAST voting stage. Ptolemaea also extends annotations by passing BLAST-only hits through strict amino acid sequence homology filters. We ran the pipeline on 700 complete ESKAPE and *E. coli* genomes, where we annotated an additional 12,363 phage defence genes that were not annotated by PADLOC or DefenseFinder. As far as we are aware, this is the only approach that provides a streamlined and reproducible means of comprehensively consolidating phage-defence annotations across multiple bioinformatic tools. Furthermore, in the cases of unmapped or conflicting annotations, it drives users to make careful considerations about the underlying biology of their gene annotations in order to complete a consensus annotation.

## Limitations

As Ptolemaea relies on external tools and databases, it inherits the limitations of its components: it can only detect systems modelled by PADLOC, DefenseFinder, or those present in the *B. cereus* group reference set, and its results will change as those resources are updated. We have not benchmarked Ptolemaea against curated ground truth, and make no claim of annotation reliability beyond what is already trusted of the component tools. Finally, consensus naming depends on a mapping between the tools’ nomenclatures, which must be maintained as they evolve. Ptolemaea is not a replacement for any one tool, but rather an aid for researchers seeking to maximise annotations rather than deciding on the ‘best’ tool to use.

## Acknowledgements

We are grateful to Elise July and Annika Gillis for validating and curating the *B. cereus* dataset, from which we derived the majority of consensus mappings, and to the creators of the defence-annotation tools PADLOC and DefenseFinder, for enabling us to further our own research. This work was supported by the project “Improved Pig Health through the Novel Application of SynBio in Phage Therapy” (2020US-IRL201), funded by the Department of Agriculture, Environment and Rural Affairs (DAERA, Northern Ireland) through the 2020 US-Ireland R&D Partnership Call. We are also grateful for use of the computing resources from the Northern Ireland High Performance Computing (NI-HPC) service funded by EPSRC (EP/T022175).

